# Gut microbiota analysis in rats with methamphetamine-induced conditioned place preference

**DOI:** 10.1101/115709

**Authors:** Tingting Ning, Xiaokang Gong, Lingling Xie, MA Baomiao

## Abstract

Methamphetamine abuse is a major public health crisis. Because accumulating evidence supports the hypothesis that the gut microbiota plays an important role in central nervous system (CNS) function, and research on the roles of the microbiome in CNS disorders holds conceivable promise for developing novel therapeutic avenues for treating CNS disorders, we sought to determine whether administration of methamphetamine leads to alterations in the intestinal microbiota. In this study, the gut microbiota profiles of rats with methamphetamine-induced conditioned place preference (CPP) were analysed through 16S rRNA gene sequencing. The faecal microbial diversity was slightly higher in the METH CPP group. The propionate-producing genus *Phascolarctobacterium* was attenuated in the METH CPP group, and the family *Ruminococcaceae* was elevated in the METH CPP group. Short chain fatty acid analysis revealed that the concentrations of propionate were decreased in the faecal matter of METH-administered rats. These findings provide direct evidence that administration of METH causes gut dysbiosis, enable a better understanding of the function of gut microbiota in the process of drug abuse, and provide a new paradigm for addiction treatment.

## IMPORTANCE

Recently methamphetamine (METH) use has become a global problem. But there is no widely accepted effective pharmacotherapeutic treatments to reduce METH abuse. Many studies explored the links between gut microbiome and brain, and what is most intriguing is that modification of gut bacteria can reverse some abnormalities of many psychiatric disorders. Therefor the use of psychobiotics may hold promise for novel treatments for METH abuse, but at first systematic researches are needed to examine whether administration of METH could cause gut dysbiotic and is there any definite relationship between specific taxa and METH administration. In this paper, we studied the gut microbiota of rats with METH-induced conditioned place preference (CPP) and very encouraging results obtained. A propionate producing genus was attenuated in the METH CPP group. Family *Ruminococcaceae* was elevated in the METH CPP group, and this family showed positive with anxiety and negative with memory in previous study.

Humans live in a co-evolutionary association with a vast ecosystem of microorganisms living within the human body, which are termed the microbiota. The adult gastrointestinal tract contains a population of 10^14^ microorganisms, a number approximately 10 times the number of human cells in the body, and the collective genome of the microbiota amounts up to 150-fold more genes than are present in the human genome^1^. Although advances in the understanding of the role of microbiota in other areas of human health have yielded intriguing results^2-5^, microbes and the brain have rarely been thought to interact. However, that belief is slowly changing^1, 6-8^. Recent investigations have rapidly uncovered a series of compelling connections between the gut and the brain^6,7,9-11^. The relationship between the brain and the gut is primarily regulated at different levels including immune, neural, and endocrine levels^12^. The gut-brain-axis comprises the biochemical signalling that occurs between the gastrointestinal tract and the CNS. Indeed, microbiotic abnormalities have been associated with conditions such as decreases in neural plasticity and neurotransmitter levels (e.g., serotonin), cognitive deficits, anxiety, depression, autism, Parkinson’s disease, and schizophrenia^8,13^.

Drug addiction, defined as compulsive, out-of-control drug use despite negative consequences, remains one of the major public health issues worldwide; hence, novel approaches are needed to augment or replace current behavioural and pharmacological interventions to treat addictive disorders^14^. The emerging links between the gut microbiome and the CNS have brought about a paradigm shift in neuroscience, with possible implications for understanding the pathophysiology and treatment of stress-related psychiatric disorders. Addiction may be viewed as a form of drug-induced neural plasticity. The current hypothesis, therefore, is that gut dysbiosis plays a key role in addictive disorders^15^.

Several possibilities exist wherein perturbations of gut microbiota might contribute to addiction. For example, the dietary and nutrition statuses of drug abusers are different from those of the general population and may in turn affect the composition of the gut microbiota. Volpe et al. have compared the composition of the intestinal flora between cocaine users and non-cocaine users and have found that cocaine users have a higher mean relative abundance of *Bacteroidetes* and a lower abundance of *Firmicutes* than nonusers^16^; are more likely to smoke; have a lower mean percentage of body fat; and consume more alcohol than nonusers. In addition, stress is one of the most significant risk factors for the development of addiction to various drugs^17^, and depression and anxiety are often comorbid with addiction. Growing evidence from studies in animals and humans indicates that gut dysbiosis can lead to altered stress responses^12, 18-22^ and depressive symptoms, each of which can contribute to addiction^7, 23-25^. What is most intriguing from a treatment perspective is that the modification of gut bacterial profiles can reverse some of the above-mentioned abnormalities^18,19,26^. More direct evidence has come from the observation that mice with fewer gut bacteria as a result of treatment with non-absorbable antibiotics show an enhanced sensitivity to cocaine reward and an enhanced sensitivity to the locomotor sensitizing effects of repeated cocaine administration^27^.

Methamphetamine (N-methyl-O-phenylisopropylamine; METH) is a commonly abused and highly addictive illicit drug. Regular use of METH is associated with a myriad of serious medical, cognitive, and psychosocial problems for drug users, which occur not only during abuse of the drug but also after cessation of use. The efficacy of the existing therapies for METH addiction, including cognitive therapy and substitution therapy, are still debated. Hence, novel approaches are needed to augment or replace current behavioural and pharmacological interventions for METH addictive disorders. The influence of the gut microbiome on the core symptoms of neuropsychiatric disorders is becoming increasingly recognized, and the microbiome might be a tractable target for novel treatment options. Therefore, the use of psychobiotics may hold great promise for novel treatments for drug addiction, but first, systematic studies are needed to examine whether administration of METH causes gut dysbiosis and whether there is a definite relationship between specific taxa and the use of METH.

In this study, a conditioned place preference (CPP) rat model was adopted, and high-throughput sequencing of 16S rRNA and bioinformatics analysis were performed to analyse the variations in gut microbiota in METH-administered rats compared with the control rats. The diversity of the bacterial community, the microbiota composition and the taxa that best characterized each group were analysed. In addition, the concentrations of propionate and butyric acid in each group were analysed.

## RESULTS

### METH-induced CPP

During the preconditioning stage of the CPP paradigm, there was no difference between the time spent in chamber A between the control group and the METH CPP group. After treatment with METH for 7 sessions, a significant CPP was established. Rats spent 374 s of their time in the METH-paired, naturally unpreferred white compartment. In contrast, saline treated-rats exhibited a natural preference for the black compartment, and no CPP was seen (Fig. 1).

**Figure 1.**
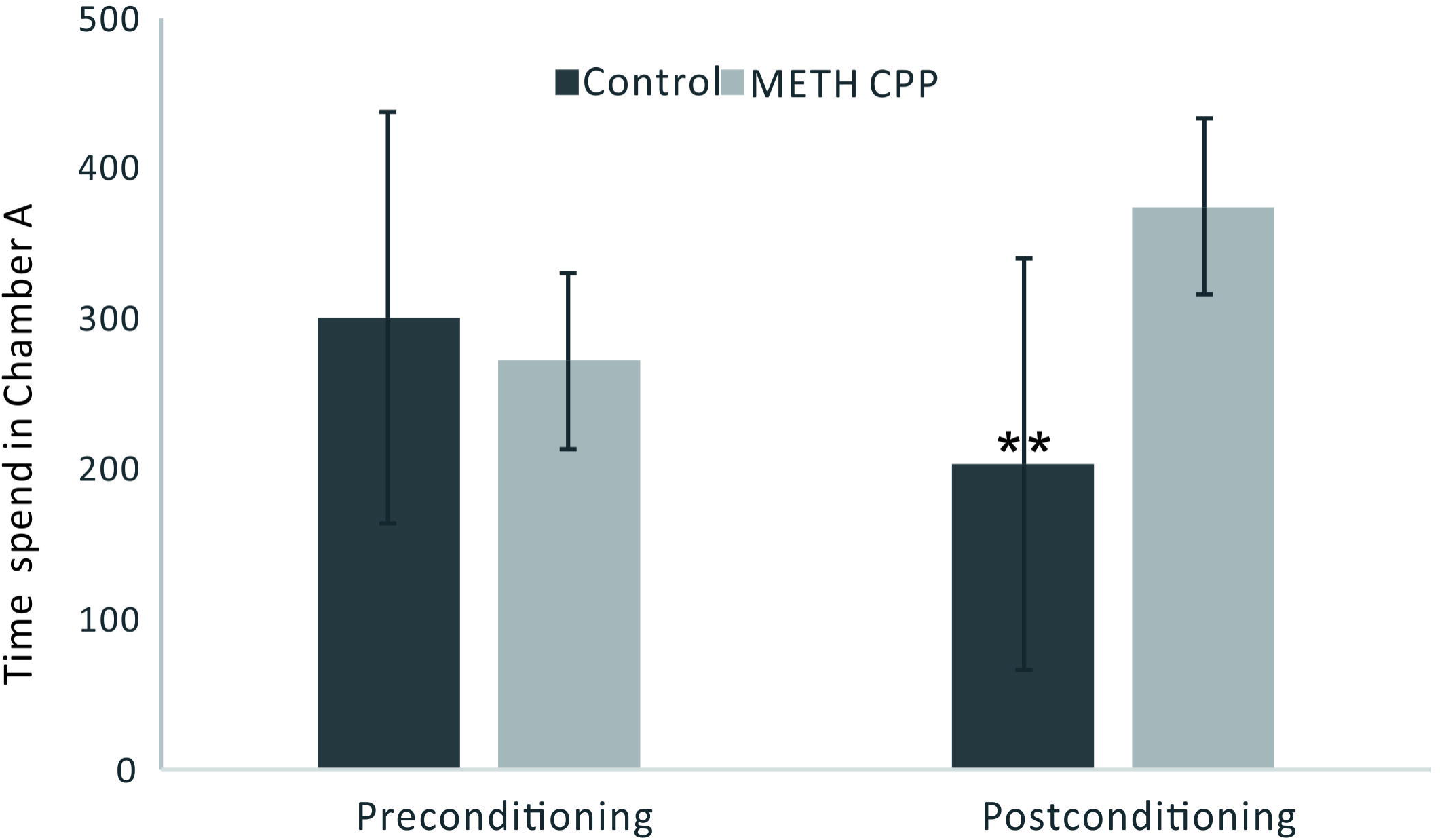
Development of CPP induced by repeated administration of METH. Data are the mean±SEM of time spent in the METH-paired chamber (Chamber A) during the CPP tests. Double asterisk indicates significant difference between the METH CPP group and the control group P < 0.01, n = 8 rats per group.

### Diversity of the microbial community

In total, approximately 318,736 sequence reads of 16S rRNA genes with an average length of approximately 439 bp were obtained after trimming and chimaera removal. A 97% similarity cut-off was used to delineate Operational Taxonomic Units (OTU)s in the downstream analyses. After subsampling, a total of 5384 OTUs were acquired (Table 1). Rarefaction curves suggested that the bacterial community was well represented because the curves became relatively flat as the number of sequences analysed increased (Fig. 2). Species richness is the number of bacterial species assigned by OTUs detected in the samples. Richness estimates were obtained from the observed number of species by extrapolation using estimators such as the ACE and Chao1 indices (Table 1). ACE and Chao1 were estimated to be 376.25/ 378.5 in the control group and 411.25/412.625 in the METH CPP group, respectively, but there were no differences between these two groups. However, the microbial diversity, indicated by the Shannon and Simpson indices, was different between the two groups. The Shannon and Simpson indices were 3.403/0.127 in the control group and 3.904/0.069 in the METH CPP group (p value for the Shannon and Simpson indices between the two groups was 0.035 and 0.038, respectively, as determined by a two-tailed t test). Because a higher Shannon and lower Simpson index indicate higher community diversity, the results suggest that the faecal microbial diversity in the METH group was somewhat greater than that in the control group. This result was unexpected, because a greater bacterial diversity is potentially beneficial to human health, but the role of bacterial diversity in CNS function remains subject to debate.

**Table 1.** Comparison of phylotype coverage and diversity estimation of the 16S rRNA gene libraries for individuals at 97% similarity from the pyrosequencing analysis

| Sample ID | Reads | 97% |  |  |  |  |  |
| --- | --- | --- | --- | --- | --- | --- | --- |
|  |  | OTU | Ace | Chao | Coverage | Shannon | Simpson |
| C 1 | 19921 | 419 | 456 | 458 | 0.997139 | 4.46 | 0.0297 |
| C 2 | 19921 | 316 | 380 | 375 | 0.996486 | 3.34 | 0.1144 |
| C 3 | 19921 | 325 | 383 | 384 | 0.996536 | 3.6 | 0.0839 |
| C 4 | 19921 | 266 | 315 | 334 | 0.997038 | 3.12 | 0.1565 |
| C 5 | 19921 | 334 | 388 | 381 | 0.996737 | 3.41 | 0.1237 |
| C 6 | 19921 | 271 | 343 | 350 | 0.996687 | 3.03 | 0.1948 |
| C 7 | 19921 | 290 | 357 | 350 | 0.996587 | 3.37 | 0.1057 |
| C 8 | 19921 | 319 | 388 | 396 | 0.996185 | 2.89 | 0.2055 |
| METH CPP 1 | 19921 | 363 | 404 | 418 | 0.996938 | 3.83 | 0.0896 |
| METH CPP 2 | 19921 | 347 | 402 | 390 | 0.996687 | 4.05 | 0.0414 |
| METH CPP 3 | 19921 | 375 | 418 | 411 | 0.996988 | 4 | 0.0505 |
| METH CPP 4 | 19921 | 304 | 362 | 367 | 0.996687 | 3.42 | 0.1071 |
| METH CPP 5 | 19921 | 362 | 414 | 424 | 0.996436 | 3.92 | 0.0491 |
| METH CPP 6 | 19921 | 363 | 436 | 435 | 0.995884 | 3.48 | 0.1465 |
| METH CPP 7 | 19921 | 438 | 490 | 492 | 0.996436 | 4.61 | 0.0228 |
| METH CPP 8 | 19921 | 292 | 364 | 364 | 0.996687 | 3.92 | 0.0475 |

Owing to significant inter-individual variation, the faecal microbiotas of the two groups could not be divided into clusters according to community composition by using unweighted UniFrac metrics and could not be separated clearly by PCoA analysis (ADONIS test, R^2^=0.08276, p value=0.191; Fig. 2).

**Figure 2.**
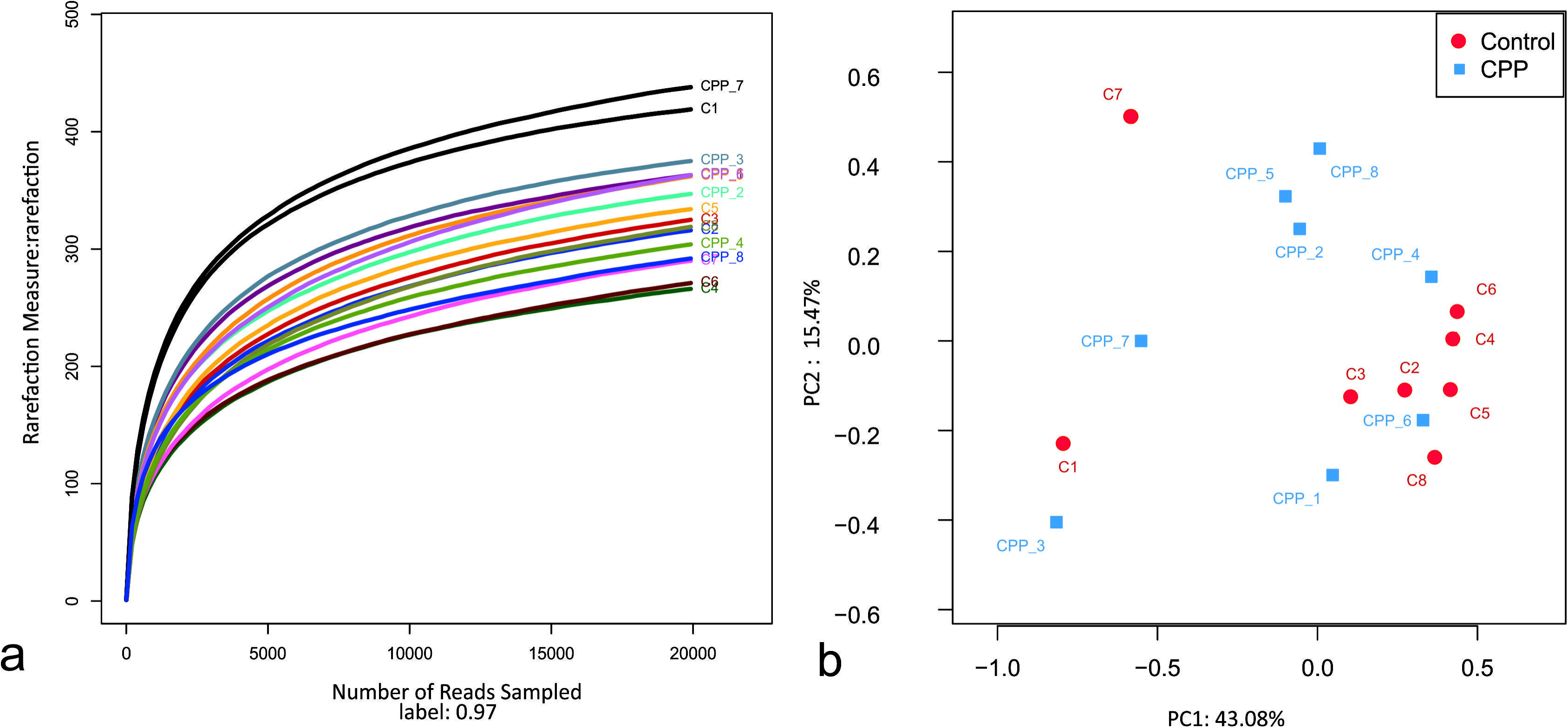
Rarefaction curve of 16S rDNA sequences of the 16 samples and principal coordinate analysis of the samples using Unweighted-UniFrac from pyrosequencing. Panel a, Rarefaction curves based on the 16S rRNA gene sequencing of the 16 samples from the METH CPP group and the control group (C represents the control group, CPP represents the METH CPP group). The rarefaction curves suggested that the bacterial community was represented well because the curves became relatively flat as the number of sequences analysed increased. Panel b, Principal coordinate analysis (PCoA) of the samples using Unweighted-UniFrac from pyrosequencing. The red dots represent the control group, and the blue squares represent the METH-CPP group. The faecal microbiotas of the two groups could not be divided into clusters according to community composition using Unweighted UniFrac metrics and could not be separated clearly by PCoA analysis (ADONIS test, R^2^=0.08276, p value=0.191).

### Altered microbiota composition in the METH CPP group

Bacteria from the sixteen samples demonstrated similar richness but different abundances. The relative bacterial community abundance on the phylum and genus levels are shown in Fig. 3. Linear discriminant analysis (LDA) effect size (LEfSe), a method for biomarker discovery, was used to determine the taxa that best characterized each population. LEfSe scores measure the consistency of differences in the relative abundance between taxa in the groups analysed with a higher score, thus indicating higher consistency. LDA revealed distinct taxa in the microbiome of the control group versus the METH CPP group (Fig. 4). *Ruminococcaceae, Bacillus, Cetobacterium, Fusobacteria,* and *Aeromonas,* which were abundant in the METH CPP group, were the key phylotypes that contributed to the difference in the intestinal microbiota composition between the METH CPP and control group. In addition, *Phascolarctobacterium,* which was more abundant in the control group, was the representative phylotype in the normal group (Fig. 5).

**Figure 3.**
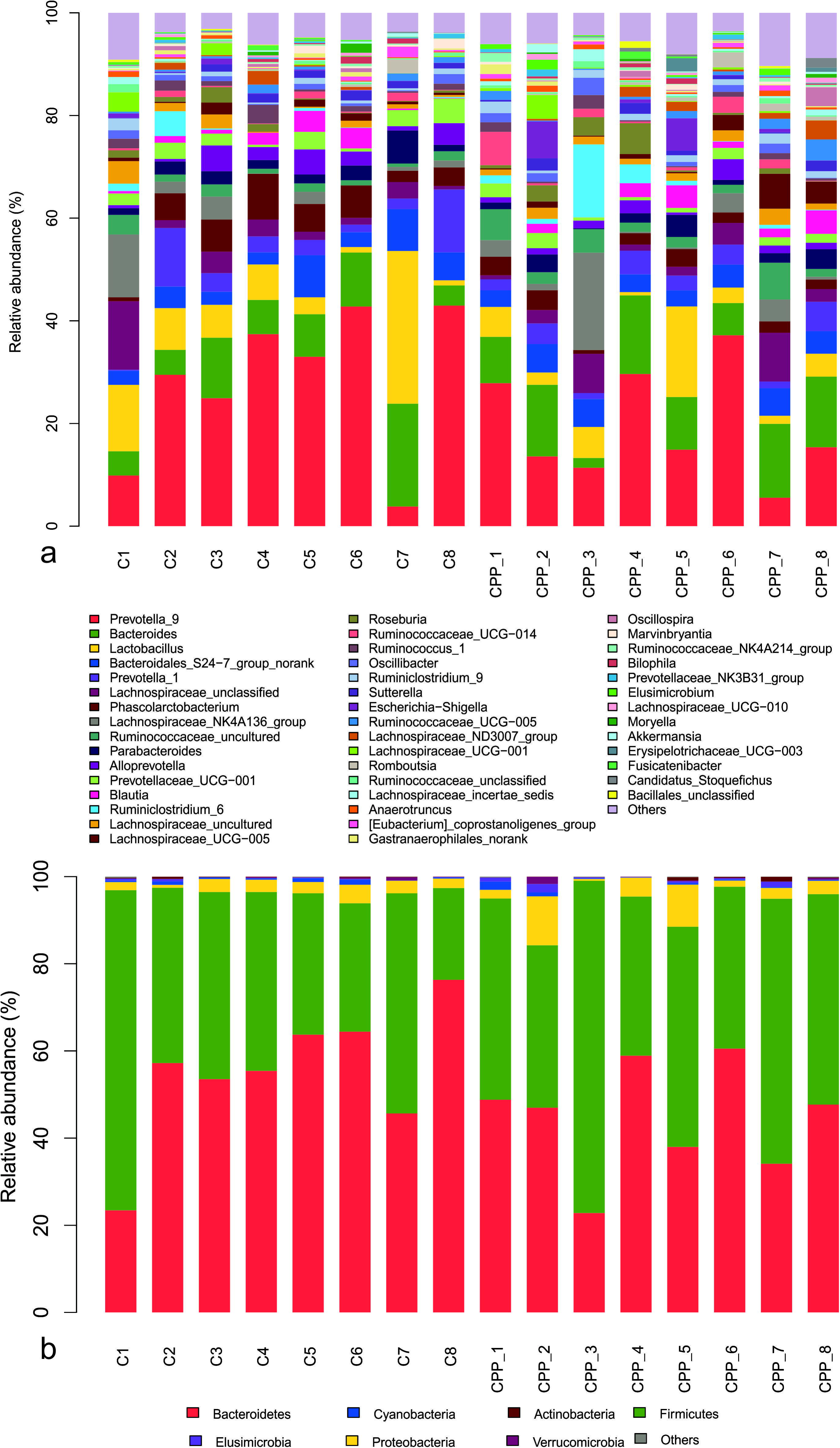
Bacterial community structures in all samples at the genus level (Panel a) and phylum level (Panel b). The abundance is presented in terms of the percentage of the total effective bacterial sequences in the sample.

**Figure 4.**
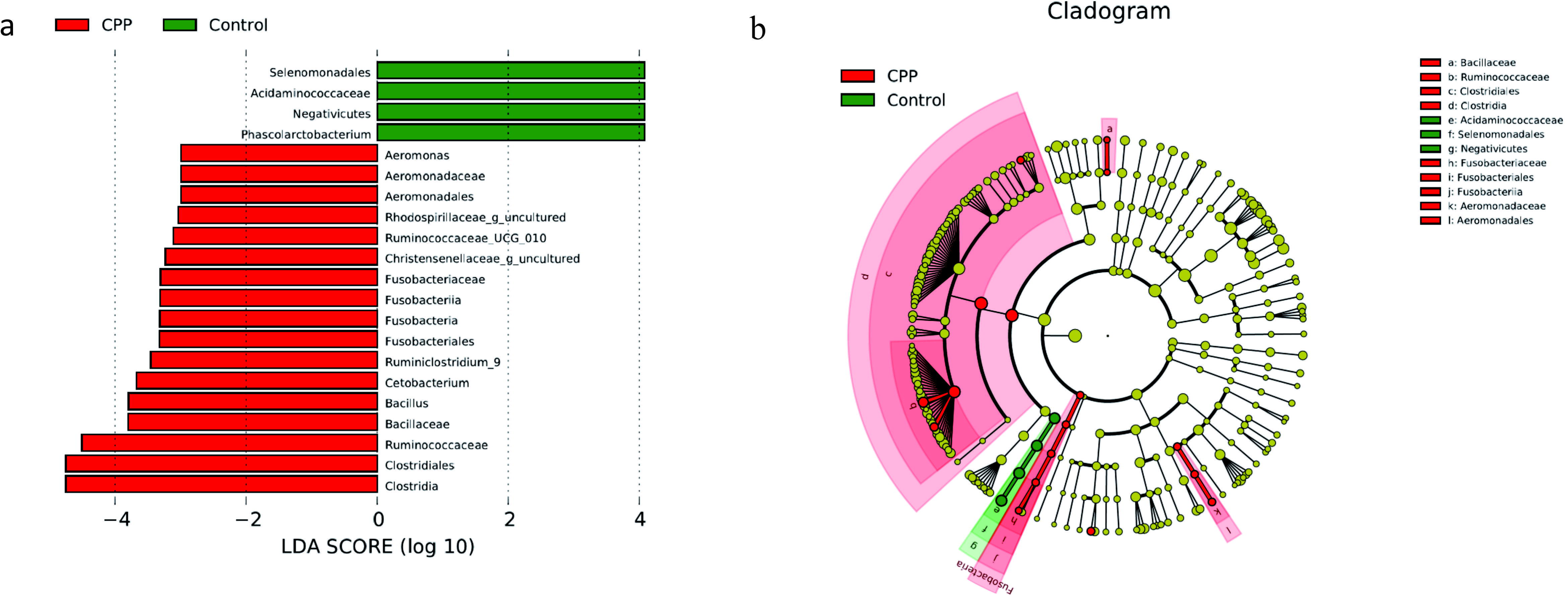
LDA shows distinct gut microbiome composition in the METH CPP group and the control group. Panel a, LDA scores, as calculated by the LEfSe of taxa differentially abundant in the two groups. Panel b, LEfSe cladogram representing differentially abundant taxa. Only taxa with LDA scores of more than 2 are presented.

**Figure 5.**
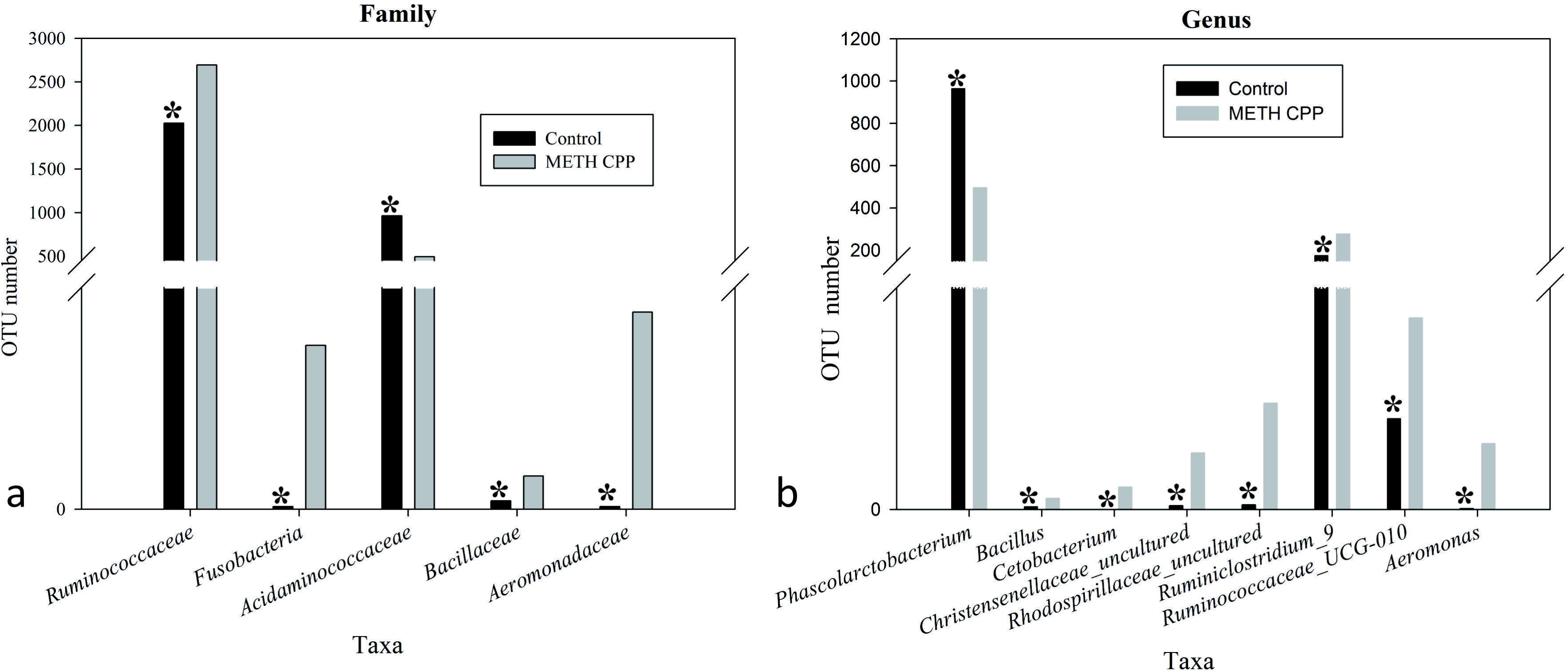
Taxonomic differences of faecal microbiota between the control and METH CPP groups. Comparison of relative abundance at the bacterial family (a) and genus (b) levels between these two groups. Student’s t test was used to determine whether differences existed between the two groups. * indicates p <0.05.

### Analysis of propionate and butyric acid

Because the genus *Phascolarctobacterium* has been characterized by producing propionate, and this genus was decreased in the METH CPP group, we proposed that propionate was also attenuated by METH administration. To test this hypothesis, we analysed the relative concentration of propionate in the faecal matter of the two groups by GC-MS. The data showed that the relative concentration of propionate in the control group was more than twice that in the METH CPP group. We also observed a trend in the variation of butyric acid, but no differences were found between these two groups (Fig. 6).

**Figure 6.**
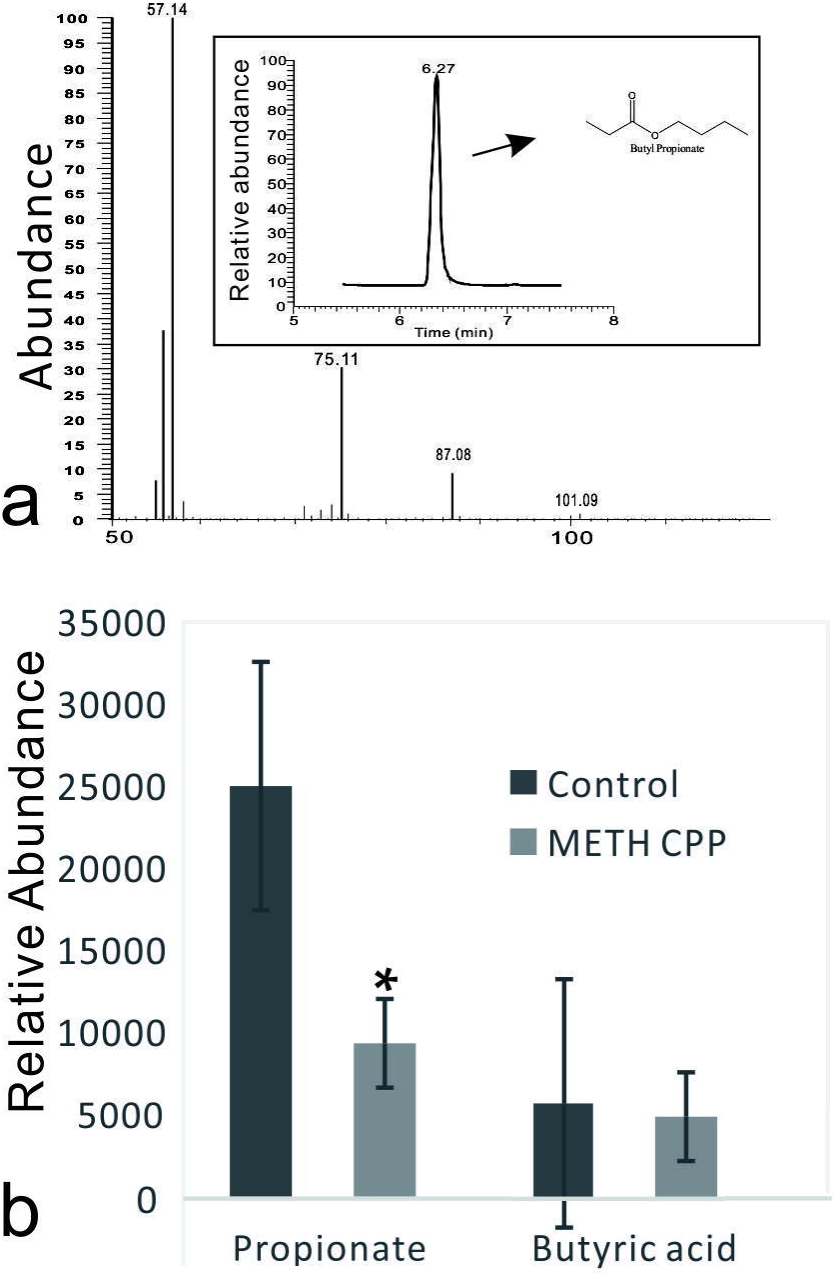
Analysis of propionate and butyric acid in the faecal matter of the control group and the METH CPP group. Panel a, Representative GC-MS spectrum of the butanol derivative of propionate in the faecal sample. Panel b, Relative abundance of propionate and butyric acid in the faecal matter of the control and METH CPP group tested through GC-MS (Means ± SEM). Each sample was analysed three times. *indicates difference between the control group and the METH CPP group (p<0.05).

## DISCUSSION

This study found differences in α-diversity in the levels of specific bacterial taxa in gut microbiomes and in the concentration of propionate associated with METH administration. To date, there has been only one study that has compared the faecal microbiota of cocaine users and nonusers and found no significant group differences in microbiota diversity^16^. In animal models of drug dependence, in contrast to studies of clinical samples, variables, including diet, living environment, age, sex, etc., can be controlled for easily; thus, the influence by factors other than drug administration can be minimized. This is the first study to explore changes in the faecal microbiota of a drug-administered animal model. In the present study, the faecal microbial diversity (estimated using the Shannon and Simpson indices) was greater in METH CPP rats. Although greater bacterial diversity is potentially beneficial to human health, its role in CNS function remains subject to debate. A previous study has demonstrated greater gut microbe diversity in active major depressive disorder (A-MDD) patients^23^. Furthermore, increased gut microbe diversity and richness have also been detected in a sample of autistic children^28^. Therefore, the precise consequences of increased bacterial diversity for METH addiction remains unclear.

In our study, intestinal dysbiosis was characterized by significant taxonomical differences between three of the major phyla in the METH CPP versus control subjects. Several taxa from *Proteobacteria* and *Fusobacteria* were significantly more abundant in METH CPP subjects, whereas the proportion of different families of *Firmicutes* showed different variation trends in these two groups. *Acidaminococcaceae* was more abundant in the control group, and *Ruminococcaceae* and *Bacillaceae* were elevated in the METH CPP group.

*Firmicutes* are the largest phylum of bacteria, comprising more than 200 genera, most of which belong to the gram-positive bacteria, in which lipoteichoic acid is a major constituent of the cell wall. The genus *Phascolarctobacterium,* belonging to the family *Acidaminococcaceae,* was the characteristic taxon in the control group. The relative abundance of this genus in the control group was approximately two-fold more than in the METH CPP group (Fig.5). This genus has frequently been reported in relationship to obesity and nutrition and had significant positive correlations with the nutritional index in humans^29^ and a high-fat diet in rats^30^. In a clinical study, the mean relative abundance of *Firmicutes* in cocaine users has been found to be lower than that in nonusers^16^; this trend was the same as that shown in body fat and the healthy eating index. Inspired by the findings suggesting a role of *Firmicutes* in obesity, we compared the weights of these two groups, but no differences were observed. Isolation of members of this genus might have been difficult because of the very narrow range of energy sources available, and this genus has been characterized on the basis of its utilization of succinate and production of propionate^31^. Propionate, a C3-short chain fatty acid, along with other short-chain fatty acids (SCFAs), small organic monocarboxylic acids with less than six carbon atoms, are major microbial metabolites produced during anaerobic fermentation in the gut^32^. SCFAs are the most important and pleiotropic functional components of microbe-to-host signalling^33^. They have shown promising effects in various diseases including obesity, diabetes and inflammatory diseases as well as neurological disorders^32,34,35^. It is now well established that SCFAs modulate colonic motility by stimulating serotonin (5-HT) secretion from gut enterochromaffin cells, which supply 5-HT to the mucosa, lumen and circulating platelets^33,36^. A recent study has found that GF mice have immature and less active microglia, an effect that can be normalized by addition of an SCFA cocktail consisting of acetate, propionate and butyrate to the drinking water^11^.

We presumed that the decrease in *Phascolarctobacterium* abundance in the METH CPP group would result in the attenuation of the concentration of propionate. To validate this hypothesis, we analysed the relative concentration of propionate in the faecal matter of the two groups by GC-MS (Fig. 6). As we expected, the concentration of propionate in the control group was more than twice that in the METH CPP group. Further studies should be carried out to validate whether serotonin release and the function of microglia are affected.

Propionate is produced in the human large intestine by microbial fermentation, and has potential health-promoting effects, including anti-lipogenic, cholesterol lowering, anti-inflammatory and anti-carcinogenic actions^37^. Although, together with acetate and butyric acid, propionate was the major short chain fatty acid produced in the gut by colonic microbiota of fermentation dietary fibres, most previous studies have focused on butyrate and to a lesser extent acetate. Consequently, the potential effects of propionic acid on physiology and pathology have been underestimated^32^. Propionate can readily cross both the gut-blood and blood-brain barriers and can have neuroactive effects, but the manner in which propionate may influence the central nervous system is unknown. Very recently and interestingly, intraventricular infusions of PA have been demonstrated to cause behavioural and brain abnormalities in rats similar to those seen in humans suffering from autism, probably via altering brain fatty acid metabolism ^38-41^. Our results thus shed new lights on the role of propionate in human health.

Two genera, *Ruminococcaceae_UCS-010* and *Ruminiclostridium_9,* which were increased in the METH CPP group belonged to the family *Ruminococcaceae*. This family displayed positive correlations with the butyrate/SCFA ratio and has been reported to be affected by different kinds of mental diseases^35^. However, we found no differences in the relative abundance of n-butyric acid between the two groups. Stressor exposure significantly changed the structure of the colonic mucosa-associated microbiota in control mice and resulted in anxiety-like behaviour, but the effect was not evident in mice fed milk oligosaccharides. Several phyla were increased in control mice compared with prebiotic-fed mice, including unclassified *Ruminococcaceae^19^*. This result indicated that *Ruminococcaceae* correlate with anxiety, which is a key characteristic of METH abusers. Short-term exposure to high-energy diets impairs memory. Rats consuming saturated fatty acids and sugar, compared with controls and rats consuming polyunsaturated fatty acids, were impaired in hippocampal dependent place recognition memory, even though all rats consumed similar amounts of energy. *Ruminococcaceae* have shown a range of macronutrient-specific negative correlations with place memory^42^; thus, this phylum plays an important role in cognitive function. In our results, the abundance of the genera *Ruminiclostridium_9* and *Ruminococcaceae_UCG_010,* both of which belong to the family *Ruminococcaceae,* was much higher in the METH CPP group than in the control group. METH abuse may result in cognitive deficits, especially in the cognitive domains of memory, attention and executive function. Therefore, the elevated abundance of *Ruminococcaceae* may be involved in the deleterious effects of METH on cognitive processes. In summary, recent research has provided strong evidence for the role of the commensal gut microbiota in brain function and behaviour, but, to date, there has been no systematic study of the involvement of gut microbiota in drug addiction. This work is the first meaningful effort to determine the relationship between METH administration and gut microbiota. The faecal microbial diversity was greater in METH CPP rats than that in the control rats, and the phylum *Firmicutes* accounted for the majority of the altered taxa. The most significant finding is that the propionate-producing genus *Phascolarctobacterium* was repressed by METH administration, and this result was further confirmed by the decrease in propionate in the faecal matter. In contrast, the family *Ruminococcaceae,* which has been reported to have a positive relationship with anxiety and a negative relationship with memory, was elevated in the METH CPP group. Although further clinical and in vivo studies are needed to better understand the mechanisms underlying the link between gut microbiota and drug administration, these findings suggest that gut microbiota manipulation might be a rational approach to treat drug addiction.

## MATERIALS AND METHODS

### Animals

Male Sprague-Dawley rats (weighing 250 g ± 20 g upon arrival; total n=16) were housed individually under controlled temperature (23 ± 2 °C) and humidity (50 ± 5%) and maintained on a 12 h/12 h light/dark cycle with free access to food and water. All animal procedures were performed in accordance with the National Institutes of Health Guide for the Care and Use of Laboratory Animals, and the experiments were approved by Jianghan University Animal Use Committee.

### Chemicals and drugs

METH (98%) was obtained from the Hubei Public Security Bureau and was dissolved in 0.9% NaCl. Propionate and butyric acid were purchased from Macklin Biochemical Technology Co., Ltd. (Shanghai, China).

### Conditioned place preference

METH-induced CPP was performed as previously described, with minor modifications^43^. The conditioned place preference system (Ningbo Anilab Software and instrument Co. Ltd, Zhejiang, China) was computer-controlled and consisted of three PVC compartments. Two identically sized conditioning compartments (L×W×H: 27 cm × 21.5 cm × 20.5 cm) had different coloured walls (white or black) and different flooring (parallel metal bars or stiff metal mesh) to provide tactile and visual cues and were separated from the neutral chamber by guillotine doors. Briefly, the CPP schedule consisted of three phases: preconditioning, conditioning, and postconditioning. The sixteen rats were randomly assigned into two groups: the control (n=8) and METH CPP group (n=8). The preconditioning phase determined the baseline preference. Rats were initially placed in the middle chamber with the doors removed for 15 min. During the conditioning phase, each rat was treated for 14 days with alternate injections of either METH (1 mg/kg, i.p.) or saline (2 mL/kg, i.p.); thus, rats were treated with METH on day 1, 3, 5,⋯, 13, and were treated with saline on day 2, 4, 6,⋯, 14. Rats were confined to the white compartment for 45 min immediately after METH administration and to the black compartment after saline injection. In the postconditioning phase, rats were re-tested for METH-conditioned place preference by being allowed free access to both the white and black chambers for 15 min. Rats in the control group received saline injections throughout the conditioning and post-conditioning phases. The time spent in each of the two compartments was automatically recorded for 15 min with a camera connected via electrical interface to a computer. Student’s t tests were used to determine whether significant place preference was established (p<0.01 was considered as a significant difference).

### Faecal sample collection and DNA extraction

Faecal sample collection was performed in metabolism cages. After the place preference test, animals were placed in the metabolism cages. The collection cup was sterilized before collection. Faecal samples were placed on ice as soon as possible and stored at -80 °C. Faecal microbial DNA was extracted from 200 mg of faeces with a QIAamp DNA Stool Mini Kit (Qiagen, Hilden, Germany) according to the manufacturer’s instructions. DNA was quantified using a NanoDrop ND-1000 spectrophotometer (Thermo Electron); integrity and size were assessed by 1.0% agarose gel electrophoresis on gels containing 0.5 mg/mL ethidium bromide. DNA was stored at -20 °C before analysis.

### High-throughput sequencing

The bacterial communities in the faecal samples were investigated by Illumina MiSeq high-throughput sequencing.

The V3 and V4 regions of the 16S rDNA gene were selected for PCR. The primers were barcoded-338F (5’-ACTCCTACGGGAGGCAGCA-3’) and 806R (5’-GGACTACHVGGGT-WTCTAAT-3’; H, W, and V were degenerate bases; H represented A, T or C; V represented G, A or C; W represented A or T), where the barcode was an eight-base sequence unique to each sample. The 20-μL PCR reaction mixture was composed of 4 μL of 5X FastPfu buffer, 2 μL of 2.5 mM dNTPs, 5 μM each of forward and reverse primers, 0.4 μL TransStart Fastpfu DNA Polymerase (TransGen Biotech, Beijing, China), and 10 ng DNA template. The following cycling parameters were used: maintain at 95 °C for 2 min, 25 cycles (95 °C for 30 s, 55 °C for 30 s, and 72 °C for 30 s), and a final extension at 72 °C for 5 min. Triplicate reaction mixtures were pooled per sample, purified using an AxyPrep DNA gel extraction kit (Axygen, CA, USA) and quantified using a QuantiFluor-ST Fluorescence quantitative system (Promega, Madison, WI, USA). Amplicons from different samples were sent out for pyrosequencing on an Illumina MiSeq platform at Shanghai Majorbio Bio-Pharm Technology Co., Ltd. (Shanghai, China). All sequences have been deposited in the GenBank Short Read Archive (SRP093459).

### Processing of sequencing data

Raw fastq files were demultiplexed and quality-filtered using QIIME (version 1.9.1) with the following criteria: (i)The 300 bp reads were truncated at any site receiving an average quality score <20 over a 50-bp sliding window, and truncated reads that were shorter than 50 bp were discarded. (ii) Exact barcode matching, two nucleotide mismatch in primer matching, and reads containing ambiguous characters were removed. (iii) Only sequences that overlapped by more than 10 bp were assembled according to their overlap sequence. Reads that could not be assembled were discarded. OTUs were clustered with a 97% similarity cutoff using UPARSE (version 7.1, http://drive5.com/uparse/), and chimeric sequences were identified and removed using UCHIME. The taxonomy of each 16S rRNA gene sequence was analysed with the RDP Classifier (http://rdp.cme.msu.edu/) against the silva (SSU123)16S rRNA database by using a confidence threshold of 70%.

### Statistical analysis

Community estimators were calculated and analysed using Mothur version v.1.30.1 (http://www.mothur.org/wiki/Schloss_SOP#Alphadiversity), including richness estimators, the Ace index and Chao1 index, and α-diversity estimators, the Simpson index and Shannon index. The rarefaction curves were generated from the observed OTUs. Principal coordinate analysis (PCoA) was used to visualize clustering patterns between samples on the basis of β diversity distances.

Microorganism features distinguishing faecal microbiota specific to METH addiction were identified using the LEfSe method (http:// huttenhower.sph.harvard.edu/lefse/) for biomarker discovery, which emphasizes both statistical significance and biological relevance. LEfSe uses the Kruskal-Wallis rank-sum test with a normalized relative abundance matrix to detect features with significantly different abundances between assigned taxa and performs LDA to estimate the effect size of each feature. An α significance level of 0.05 and an effect-size threshold of 2 were used for all biomarkers.

### GC-MS analysis of the relative concentration of propionate and butyric acid in the faecal matter

The extraction procedure of short chain fatty acids was performed as previously described^44^. Briefly, a 400-mg stool sample was used to extract short chain fatty acids using 2 mL water. The short chain fatty acids extracted by water were lyophilized and then estered by adding 500 μL 5% sulphuric acid / butanol at 90°C for 2 h. Then, the esterification ‘was stopped by addition of 500 μL 0.9% NaCl. Derivatives of short chain fatty acids were then extracted with 500 μL n-hexane and analysed with GC-MS.

GC-MS analysis of fatty acid esters dissolved in hexane phase was performed and quantified using a TRACE Ultra gas chromatograph connected to a TSQ Quantum XLS triple quadrupole mass spectrometer (GC-MS, Thermo Scientific Inc., San Jose, CA, USA). The hexane phase was analysed on a TRACE TR-5MS (30 m × 0.25 mm × 0.25 um), after a 1-μL injection with an auto-sampler (AI/AS 3000). Injections were performed in split mode with a split ratio of 20:1. Helium was used as the carrier gas, with a flow rate of 1 mL/min. The inject inlet and ion source temperatures were 220 °C and 240 °C, respectively. The temperature sequence was programmed as follows: 60 °C as an initial temperature for 6 min, and then a 4 °C/min ramp to 120 °C, held at 120 °C for 10 min. Each sample was tested three times. Student’s t test was used to determine whether difference existed between the two groups (p< 0.05).

## ACKNOWLEDGEMENTS

This study was supported by the project of the Education Department of Hubei Province (B2016300). We thank Professor Chaoying Li in the Wuhan Institute of Biomedical Science and Jianghan University for help in proof-reading the article. We thank Dongmei Huang in Majorbio Bio-Pharm Technology Co. Ltd for help in data analysis.

## AUTHOR CONTRIBUTION

This study was conceived and designed by T.T.N. The experiments were conducted by T.T.N and X.K.G. The data analysis was done by T.T.N. L.L.X did the GC-MS analysis of propionate in the faecal matter. B.M.M did the DNA extraction work. All authors reviewed the manuscript.

